# Showcasing the Rhizosphere of *Medicago Polymorpha’s* Ability to Produce Biofuels, Fix Nitrogen, and Demonstrate Similarities to Native Plant Species During a Drought Season

**DOI:** 10.1101/2025.06.21.660891

**Authors:** Ruben Munoz, Natasha Ojano, Beatriz Paz, Sabrina Razaqi

**Affiliations:** UCLA

**Keywords:** *Medicago polymorpha*, rhizosphere, metabolic activity, agriculture, drought

## Abstract

Acknowledging the impact of environmental change on microbial communities, provided the distinct rationale to the role of a plant’s capabilities within its underground network of life. Wildfires are associated with a significant decrease in microbial enzymatic activity within days, yet the long-term effects remain understudied. By focusing on microbial modifications, influenced by drought and shifts in community and functional profiles, this study contributes to understanding relationships in plant species. This study investigated the microbial composition and functional characteristics of the bacteria isolated from the rhizosphere of *Medicago polymorpha* during the wildfire season at Sage Hill. It aimed to assess alterations in microbial life post-drought, honing on *Actinobacteria*, *Proteobacteria*, and *Firmicutes*, the most abundant and climate-tolerant phyla. Key results showed bacterial isolates capable of cycling biofuels and nitrogen, surprisingly resembling native species more than invasive, and showing similarities to the past wet season than to the current arid one. Statistical analysis revealed significantly different bacterial abundances and identities compared to the plants present within the same season, highlighting *Actinobacteria* and *Proteobacteria* as highly responsive to climate changes. Further analysis showed greater similarities to native species despite *Medicago polymorpha’s* invasiveness. Over half of the microorganisms hydrolyzed their environment and produced biofuels. One-third generated siderophores and 60% demonstrated nitrogen fixation. The results emphasize the understanding of microbial shifts that occur in plant species, using the new soil site of *Medicago polymorpha* as a model of how arid ecosystems inform agricultural development. These findings support microbial resilience and provide strategic agriculture in drought-prone environments.

## Introduction

During the first few months of 2025, Los Angeles experienced unusual weather conditions for the time of year, causing wildfires to spread through the Palisades and Pasadena (Stelloh et al., 2025). Dry, drought-conducive conditions are the result of climate change, resulting in wildfires and other extreme weather conditions (Climate.gov, 2025). These unusual events informed research into the potential impacts of dry soil on microbial life due to the historic drought California continues to experience. In recent years, rising temperatures and population growth with increased “longevity and severity of climate change” droughts caused a need for water reservations in urban California landscapes (Lee, Nemati, Dinar, 2022). Climate change is significantly impacting water preservation, which has become an economic concern for California agriculture and microbial communities as the rhizosphere of plants are becoming abnormally dry (Stone & Johnson, 2022).

The soil for this research was collected from Sage Hill, located on UCLA campus and next to an invasive plant named *Medicago polymorpha*. Known also as a California burclover, *Medicago polymorpha* is an invasive plant near roadsides or vacant lots, and also seen in the Mediterranean (CAL-IPC). However, it is also known for its tolerance in drought conditions, as it can adapt with environmental changes (USDA-NRCS).

The significance of this project is to research drought-resistant bacteria and its impact on microbial communities and if the recent fires have had any impact. As fires are both destructive and essential, they affect diversity such as native flora and invasive species (Thapa et al., 2023). Understanding impacts of fires and drought on ecosystems and bacterial communities is important for our project to have a deeper understanding on effects these conditions cause in the rhizosphere of Medicago. While the fires burn biomass and create new carbon sources for both the bacteria and remaining flora called “black carbon,” it has both consequences and benefits to the bacteria and native environment in the short and long term (González-Pérez, José A et al., 2004). Bacterial sensitivity and activity to the fires depends on the duration, temperature, and water content in the soil (Barreiro and Díaz-Raviña, 2021). In other words, as these fires are causing destruction, the dryness of soil is the biggest cause for wildfire spread. Overall, it inspired research of the impacts of drought on microbes and finding ways to improve microbial diversity and quantity, as well as how to improve agriculture to rely less on fertilizers. This research idea led to our research question: How does drought impact microbial composition and function at Sage Hill in the rhizosphere of *Medicago polymorpha*?

With this, the overall hypothesis is since drought conditions reduce microbial activity and soil moisture, while surviving drought and fire-resistant microbes, if the rhizosphere of *Medicago polymorpha* is impacted by drought, then there will be a decrease in overall microbial activity, like slower growth and metabolism, and an increase in relative abundance of Proteobacteria, Actinobacteria, and Firmicutes in drought affected areas.

Therefore, the goal for this research project is to find drought-resistant bacteria and how drought affects microbial communities. As a result of these droughts, it led to wildfires and other causes within microbial communities. The overall goal is finding drought-related bacteria between the rhizosphere of a bacteria impacted and non impacted by the drought. With the help of CAL eDNA, who received a sample of our soil to analyze, we concluded a few drought-related bacteria were present (Meyer et al., 2019). As well as conducting experimental strategies such as functional assays: siderophore, carbohydrate, and nitrogen fixation and biochemical assays: catalase and oxidase, both assays have some growth and positive results. These assays and CALeDNA are a key step that have given answers to our research question of the impact drought has on microbial function, composition, and interpreting our drought and fire-resistant soil.

Evidence has shown many answers to our hypothesis and data about our soil and plant, *Medicago polymorpha*. Our data through CAL eDNA and assays have provided a deeper understanding of our plant and drought-related bacteria present. With this, a comparison of our and past cohort(s) data will be important for analysis to provide a deeper understanding of our drought-related soil vs non or wet soil within the same area. Overall, this research is not only to understand how drought can impact microbial communities, but how to improve agriculture and rely less on fertilizers, and if any of the recent fires have had any impact on the bacteria and soil.

## Materials and Methods

### Soil Enrichment and Cultivation by Serial Dilutions

Soil was collected from the rhizosphere of *Medicago polymorpha* by the instructional team. The sample was sifted to remove large debris and prepared for serial dilution. Serial dilutions were performed using a portion of the collected soil. Diluted samples were plated on agar using glass beads to ensure even distribution, then incubated at 30 °C until colonies developed.

### Isolate Purification

Distinct colonies were selected from the initial plates and streaked individually on *Rhizobium* defined medium (RDM) and International *Streptomyces* Project medium 4 (ISP4) for purification. Each isolate was incubated at 30 °C for 24–28 hours. Subsequent rounds of streaking and incubation were performed to ensure purity.

### Gram Staining

To assess isolate purity and cellular morphology, Gram staining was performed. Colonies were mixed with 4-5 µL sterile water on a microscope slide and air-dried. All slides were processed using an automatic Gram stain machine. Microscopy was performed using a bright-field microscope.

### Soil Characterization

#### Soil pH

3 mL of sieved soil was diluted with sterile water to a final volume of 9 mL (1:3 dilution), mixed for 30 minutes, and allowed to settle. Two pH strips were used to assess the final pH.

#### Water Content

Two aluminum dishes containing 20-30 g of soil were dried at 105 °C for at least 24 hours, cooled in a desiccator, and reweighed. Drying continued until weight change was within 5%. Gravimetric moisture content was calculated: θg = (m – d)/d x 100.

#### Active Carbon

A standard curve was prepared using KMnO_4_ solutions (0.005 M-0.02 M) adjusted to pH 7.2. Absorbance at 550 nm was measured. For testing, 5 g of air-dried soil was mixed with 20 mL 0.02 M KMnO_4_ and shaken. After settling, 500 µL supernatant was diluted, and 5 mL was measured spectrophotometrically. Active carbon was calculated using: Active C (mg kg^−1^) = [0.02 mol/L – (a + b × absorbance)] × (9000 mg C/mol) × (0.021 L / 0.005 kg).

#### NPK Testing

Commercial NPK kits were used. For nitrogen, soil was mixed with Nitrate Extract and Nitrogen Indicator; results were read based on color change. Phosphorus tests involved adding extract and an indicator followed by a tablet; results were determined by blue color intensity. Potassium tests involved titration with an indicator solution to a blue endpoint.

### Functional Assays

#### Siderophore Production

Chromeazurol S (CAS) assay overlays were used to detect siderophore production. Isolates and positive control (*E. coli*) were grown on RDM and ISP-4 plates. Growth was then picked off and the plates were overlaid with CAS media. Color change from blue to yellow indicated siderophore production.

#### Carbohydrate Polymer Degradation

Tryptone-yeast carboxymethyl cellulose (TY-CMC) plates were inoculated with isolates and a *Bacillus safensis* control. Sets of plates were incubated at 30 °C for 3-7 days depending on the triplicate. After incubation, plates were flooded sequentially with Congo red (15 min), 1 M NaCl, and 1 M HCl to visualize zones of clearing.

#### Nitrogen Fixation

Isolates were initially plated on N2-BAP to confirm growth. Then, DF media tubes were stab-inoculated with isolates and *Burkholderia unamae* control and incubated at 30 °C. Cone-shaped growth indicated nitrogen fixation. Isolates showing growth were further tested on JMV media using the same procedure.

### Biochemical Assays

#### Catalase Test

Gram-positive isolates were tested for catalase activity. A loopful of culture was placed on a slide with hydrogen peroxide; bubbling indicated a positive reaction.

#### Oxidase Test

Colonies were applied to oxidase test paper using a wooden stick. A positive result was a blue or purplish-black color within one minute.

## Statistical Analysis

### Metadata File Preparation for Cultivation-Independent Analysis

Metadata files were prepared to support cultivation-independent microbial community analysis. Data was organized using Excel, with each row representing a unique soil sample and each column corresponding to a specific environmental or collection parameter. It was ensured that sample names were exact matches with sample identifiers used in the OTU table. Columns contained either categorical or numerical data. Once formatted, the metadata file was saved as a .txt file. Preliminary Community Composition Analysis Using Pivot Tables

An initial cultivation-independent analysis was performed using Excel. The ASV table provided by the instructional team was imported into the program. A Pivot Table was inserted with taxonomic ranks under “ROWS” and sample site data under “VALUES.” “Sum” was used in the Value Field Settings to reflect the abundance of total sequences. Eukaryota and Fungi were filtered out. Visual charts (stacked bar, donut, etc.) were generated to visualize these microbial community profiles.

### Statistical Analysis of Metagenomic Profiles (STAMP)

STAMP was used to analyze taxonomic and functional profiles to generate PCA plots, bar graphs, and extended error graphs. Statistically significant results were determined with a p-value threshold of <0.05. Both metadata and ASV profile files, saved as txt. files, were downloaded, modified, and uploaded to the STAMP program and set to analyze selected phylum level. Comparisons were conducted with either multiple groups, two groups, or two samples. Filters and various statistical properties were applied to analyze relevant data. The graphs were then saved as high-resolution images at 300 DPI.

### 16S DNA Trimming and Preparation for Cultivation-Dependent Analysis

16S rDNA was extracted from purified isolates from Spring ‘24 109AL quarter, through lysis and PCR. PCR was done for 35 cycles in a three-step repeated cycle of denature at 98°C for 30 seconds, anneal at 55°C for 10 seconds, and extended 30°C for 10 seconds. Once the cycle was finished it does a final extension at 72°C for 5 minutes. Gel electrophoresis verified successful 16S rDNA extraction with a band at ~1,500 bp. The 16S nucleotide sequences were trimmed by a max of 150bp on both ends with MEGA12. The trimmed file was saved as a txt. file. The structure of the 16S rDNA was verified by pasting the trimmed rDNA sequence into RNA 2D templates (R2DT).

### 16S DNA Nucleotide BLAST

The trimmed txt. file 16S DNA sequences were obtained and NCBI BLAST was used to input the sequence to identify the 16S information. The top 3-4 hits were downloaded and those sequences were saved to build phylogenetic trees as FASTA files. Good hits had high total scores, high max scores, query coverage greater than 97%, and E-values < 1×10^−6.

### 16S Nucleotide Alignment and Phylogenetic Tree

The following files were used to build a phylogenetic tree; top 3-4 hits BLAST sequences FASTA file, the original 16S isolate DNA, and an additional outgroup (E. Coli). MEGA12 was used to align the sequences, delete unaligned base pairs, build the phylogenetic tree, observe for bootstrap values greater than 75, and adjust the phylogenetic tree accordingly to make a reliable tree.

## Results

### Soil Characteristics and Chemical Composition

The expectation was to see low moisture and nutrient availability as well as an acidic environment due to the drought conditions. The soil sample collected on Sage Hill from the rhizosphere of *Medicago polymorpha* (KOBO K3008-T9) was characterized based on soil moisture content, mineral composition, soil pH, and active carbon within the rhizosphere. After two rounds of drying, the dry weight analysis averaged at 4.385% soil moisture, as shown in Table 2. Additionally, the pH of the soil was determined to be 6, indicating a slightly acidic environment in comparison to a neutral level of 7. The soil sample mineral composition was analyzed through a NPK soil test kit, assessing the levels of nitrogen, phosphorus, and potassium. The test kit revealed relatively low levels of nitrogen (0-30 lb/acre) and potassium(1-120 lb/acre), depicted in Table 3. To determine the active carbon from the soil sample a standard curve of potassium permanganate was utilized. The absorbance in wavelength was measured at 0.275 nanometers and the active carbon equated to 581.78358 mg kg^−1^.

**Table 1.** Summary of Isolates Table. This table presents a summary of the status, macro-morphology, micro-morphology, key functional assays, and isolate identity hypotheses of all pure isolates.

| Isolate Name | Isolate Status | Macro-morphology | Micro-morphology | Key Functional Assay |
| --- | --- | --- | --- | --- |
| W25109<br>CMM23<br>0R03 | Negative | Orange, circular, flat | G+, Bacilli with spores | Siderophore(-),<br>Carbohydrate (-),<br>Nitrogen N2-BAP(+) DF(+)<br>JMV(+) |
| W25109<br>CMM23<br>0R06 | Negative | Translucent/yellow, circular, flat | G-, Coccus | Siderophore(-),<br>Carbohydrate(+),<br>Nitrogen N2-BAP(-) DF(NA)<br>JMV(NA) |
| W25109<br>CMM23<br>0R09 | Negative | Pale yellow,<br>circular, flat | G-, Bacilli<br>with spores | Siderophore(-),<br>Carbohydrate (+),<br>Nitrogen N2-BAP(+) DF(+)<br>JMV(+) |
| W25109<br>CMM23<br>0R16 | Negative | Pale yellow,<br>circular, flat | G-, bacillus | Siderophore(-),<br>Carbohydrate (+),<br>Nitrogen N2-BAP(+) DF(-)<br>JMV(+) |
| W25109<br>CMM23<br>0R18 | Negative | Pale yellow,<br>circular, flat | G-, bacillus<br>with spores | Siderophore(-),<br>Carbohydrate (+),<br>Nitrogen N2-BAP(+) DF(+)<br>JMV(+) |
| W25109<br>CMM23<br>0R20 | Negative | White, circular,<br>flat | G-, bacilli | Siderophore(+)Carbohydrat<br>e (+),<br>Nitrogen N2-BAP(+) DF(-)<br>JMV(+) |
| W25109<br>CMM23<br>0I01 | Negative | Yellow, circular,<br>flat | G-, bacilli | Siderophore(+)<br>Carbohydrate (-),<br>Nitrogen N2-BAP(-) DF(NA)<br>JMV(NA) |
| W25109<br>CMM23<br>0I02 | Negative | Yellow, circular,<br>flat | G-, bacilli | Siderophore(+), Carbohydrate (-),<br>Nitrogen N2-BAP(-) DF(NA)<br>JMV(NA) |
| W25109<br>CMM23<br>0I03 | Negative | White, circular,<br>flat | G+, bacillus<br>with spores | Siderophore(-),<br>Carbohydrate (+),<br>Nitrogen N2-BAP(-) DF(NA)<br>JMV(NA) |
| W25109<br>CMM23<br>0I04 | Negative | White, circular,<br>flat | G-, bacillus<br>with spores | Siderophore(-),<br>Carbohydrate (+),<br>Nitrogen N2-BAP(+) DF(-)<br>JMV(-) |
| W25109<br>CMM23<br>0I06 | Negative | Pale yellow, flat,<br>circular | G+, bacilli | Siderophore(-),<br>Carbohydrate (+)<br>Nitrogen N2-BAP(+) DF(+)<br>JMV(+) |
| W25109<br>CMM23<br>0I11 | Negative | White, circular,<br>flat | G+,<br>streptococcus | Siderophore(-),<br>Carbohydrate (-)<br>Nitrogen N2-BAP(-) DF(NA)<br>JMV(NA) |
| W25109<br>CMM23<br>OI14 | Negative | Yellow-orange,<br>circular, flat | G-, bacilli | Siderophore(+)Carbohydrate (-)<br>Nitrogen N2-BAP(+) DF(+)<br>JMV(+) |
| W25109<br>CMM23<br>OI19 | Positive<br>16S<br>DNA | Red/pink,<br>circular, flat | G-,<br>pleomorphic | Siderophore(-),<br>Carbohydrate (-),<br>Nitrogen N2-BAP(-) DF(NA)<br>JMV(NA) |
| W25109<br>CMM23<br>OI20 | Negative | Tan-brown,<br>circular, flat | G+,<br>streptococcus | Siderophore(+)Carbohydrate (+),<br>Nitrogen N2-BAP(+) DF(+)<br>JMV(+) |

**Table 2.**
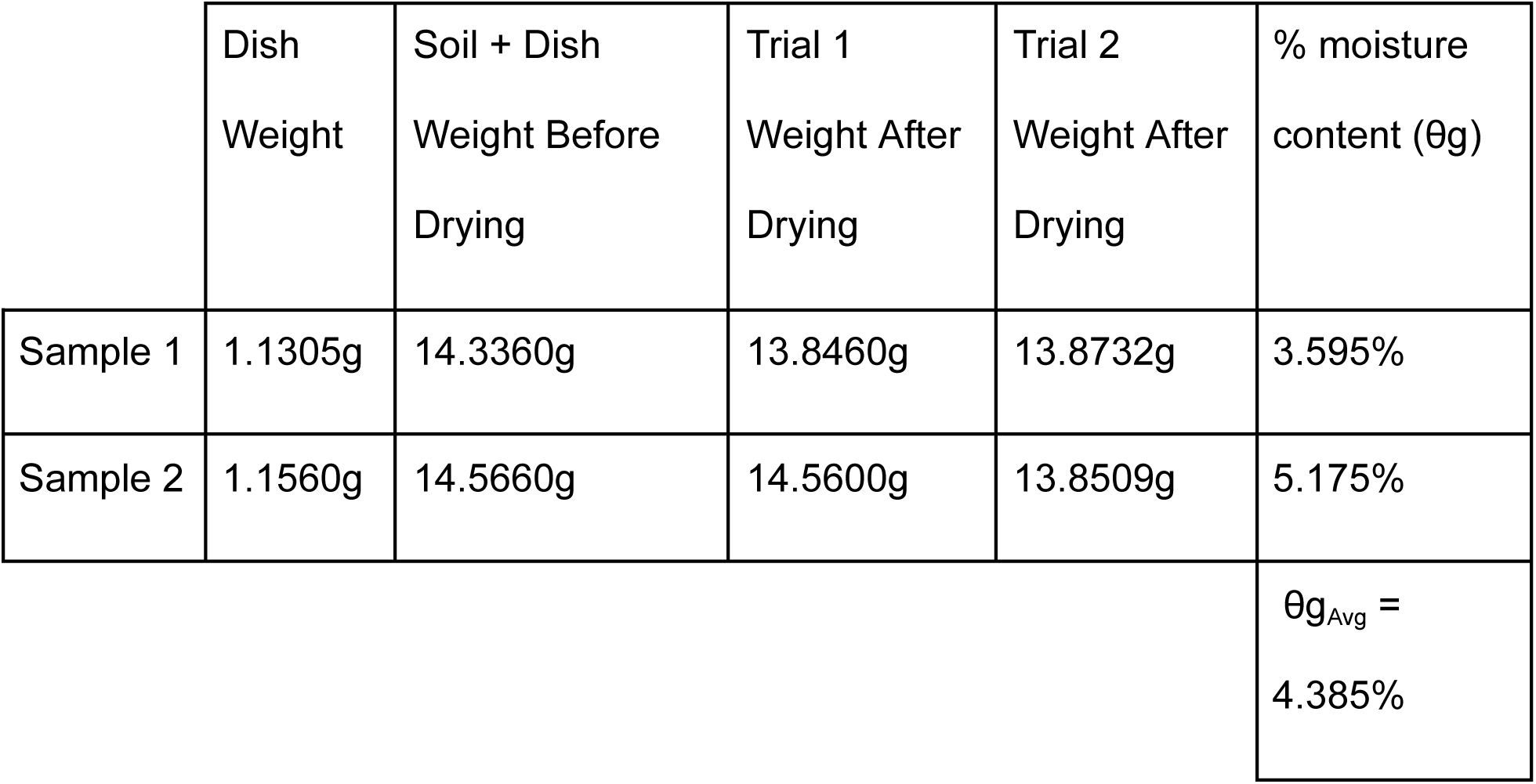
Calculations of the Soil Moisture Content The table depicts the weights for each soil sample taken before and after the drying 24 hour drying periods. Each sample represents the percentage of water content within the soil and averaged between the two samples.

**Table 3.**
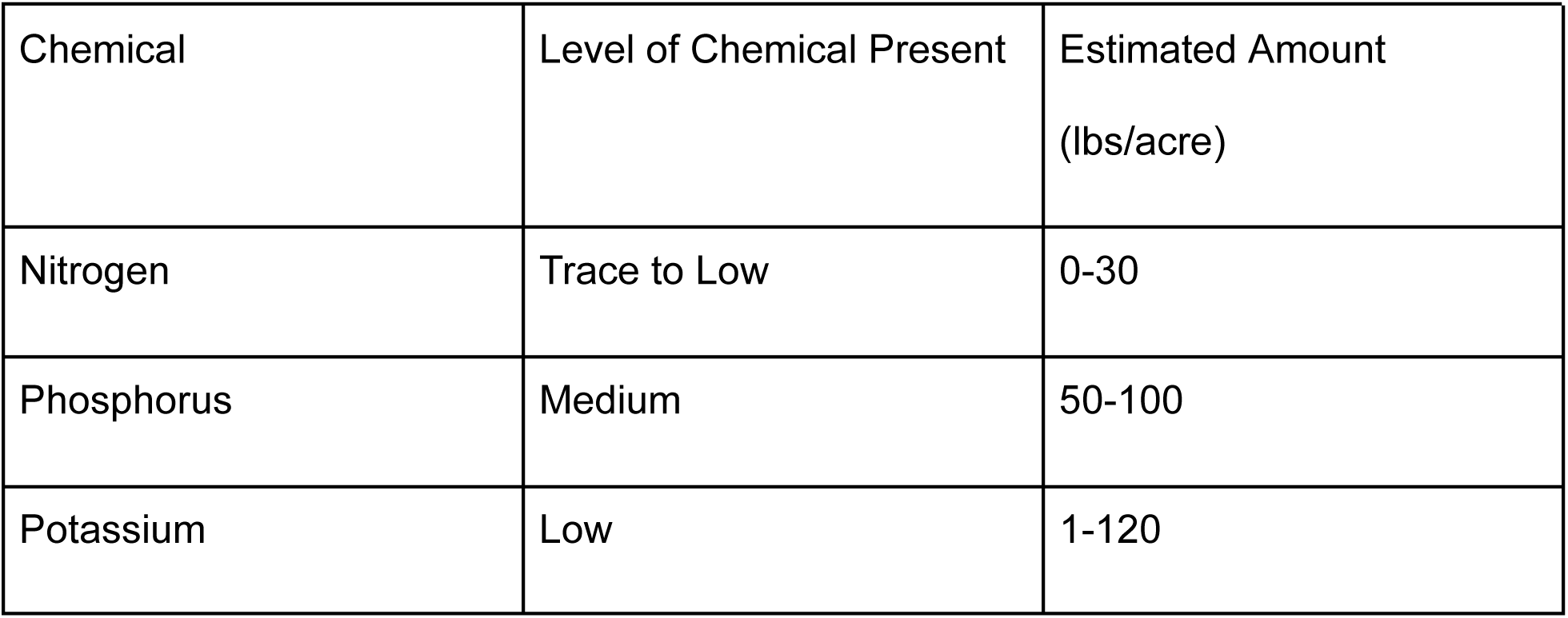
| Numerical Findings of the NPK Soil Test The presence of each chemical was measured according to their own scaling levels in pounds per acre based on the NPK Soil Test Kit provided. Nitrogen and Potassium were measured at relatively lower levels than their standard, representing a significant change in chemical existence.

| Chemical | Level of Chemical Present | Estimated Amount<br>(lbs/acre) |
| --- | --- | --- |
| Nitrogen | Trace to Low | 0-30 |
| Phosphorus | Medium | 50-100 |
| Potassium | Low | 1-120 |

These soil characteristics provide information about the environment that the bacteria survived in and how they’ve adapted to it which may alter their activity. According to Zhong et al. (2010), soil biodiversity is influenced by pH, nitrogen, potassium, and phosphorus and has consequences to bacterial survival. Given the low soil moisture content and mineral composition within an acidic environment, this may influence bacterial composition and variation.

### Species Identification Process

Through serial dilutions and three rounds of purification, the pure isolates were used to further analyze the bacterial species present. Gram staining enhances the ability to identify characteristics of the pure colonies such as, uniformity and microbe classification based on cell morphology. The analysis of the physical characteristics determined the experimental pathways taken to assist in bacterial identification. Using the automated gram stainer, 15 out of 40 isolates were confirmed to be pure with their given morphology, as shown in Table 1. Photos of both gram positive and negative bacteria were taken at 100X magnification and identified in their corresponding bacterial composition, represented in **Figures 1B** and **1C**.

**Figure 1.**
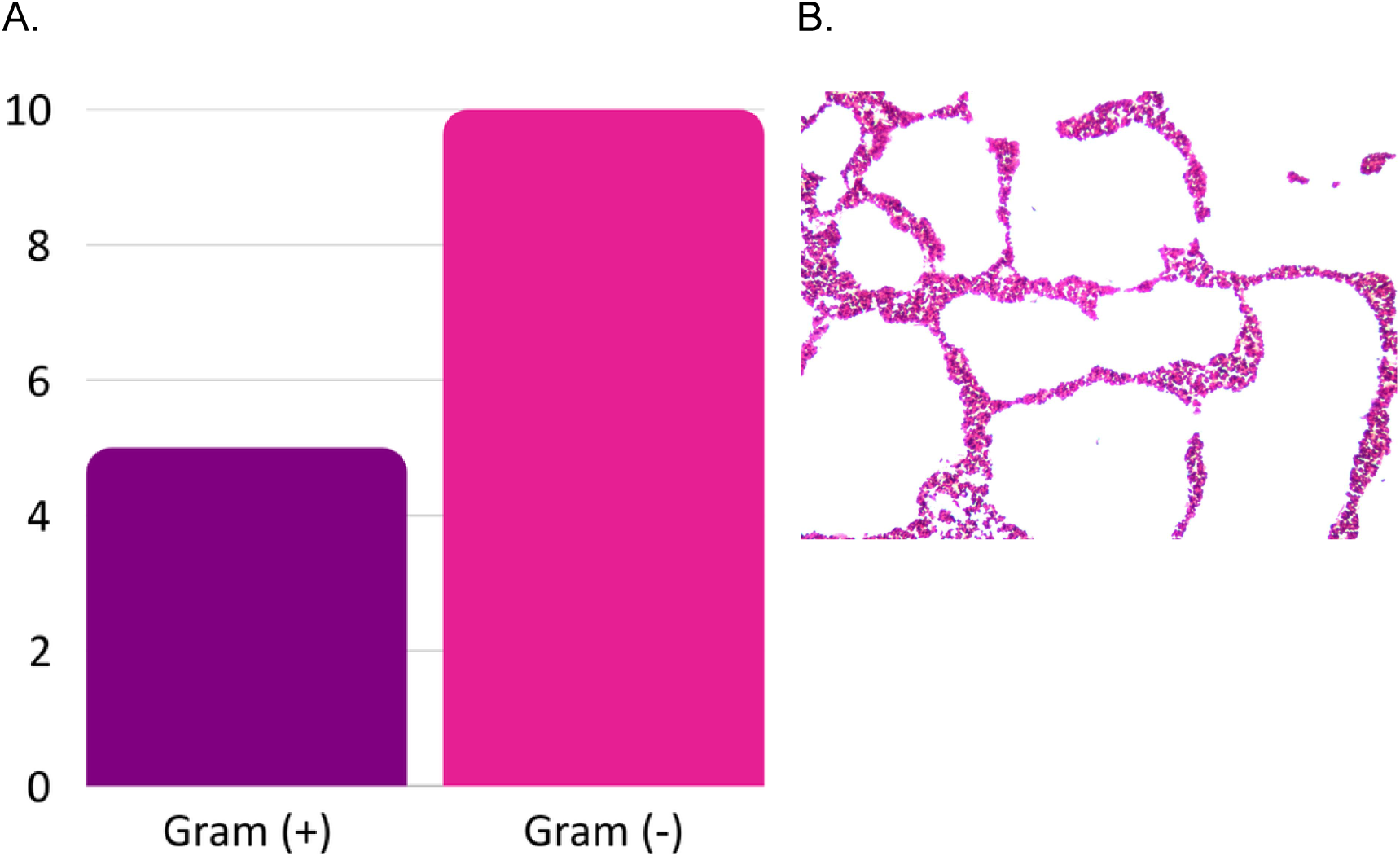
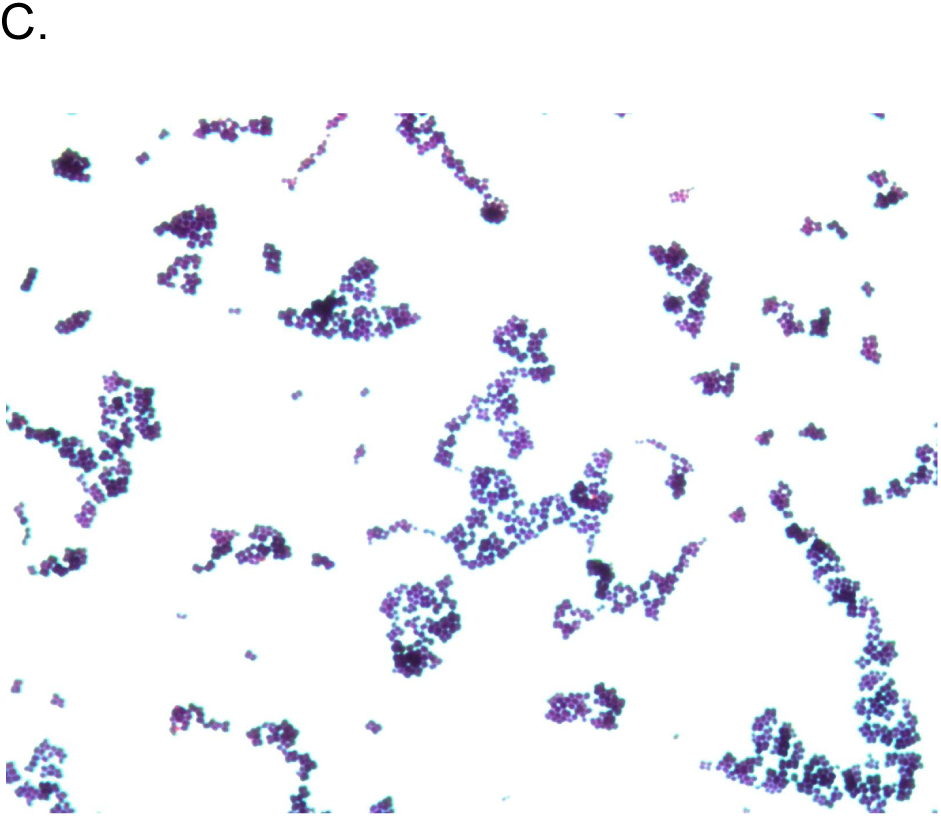
Gram Staining of Pure Isolates. **A)** After performing gram staining on all isolates, 15 were determined to be pure, with 5 being gram (+) and 10 being gram (-). **B)** Isolate W25109CMM230R09 was determined to be gram (-). It had bacilli morphology with spores. No measurements were taken, so these results were omitted. **C)** Isolate W25109CMM230I11 was determined to be gram (+). It had streptococci morphology. No measurements were taken, so these results were omitted.

## Functional Analysis

### Bacterial Isolates Associated with Siderophores

Due to siderophores being an energy-expensive molecule for microorganisms to produce, they would be less active to their community and reduce expression of secondary metabolites (Breitkreuz *et al*. 2021). Siderophores are important because the molecules are produced by microbes to scavenge iron, which is a crucial nutrient for plant growth and present in soil (Wang et al. 2022). As shown in **Figure 2C** and **2D**, two trials were conducted on the 15 isolates which demonstrated that the majority of the bacteria tested failed to locate and transport iron, signifying an iron-limited environment. In both trials, 5 of the isolates produced siderophore, and 10 isolates didn’t produce siderophore, displayed in Table 1. In **Figure 2A**, a positive result was shown when the CAS overlay was poured onto the plates showing a color change from blue to yellow indicating production of siderophore. In Figure 2B, a negative result was shown when the CAS overlay was poured onto the plate resulting in no color change as the plate stays blue. The results indicate the majority of the bacterial isolates were not capable of producing siderophores; indicating a reduction of the energy-expensive molecule allows bacteria to survive under drought conditions. As siderophores are energy-expensive they thrive in faster growing environments, however, due to the drought-related conditions present these microorganisms are slower growing.

**Figure 2.**
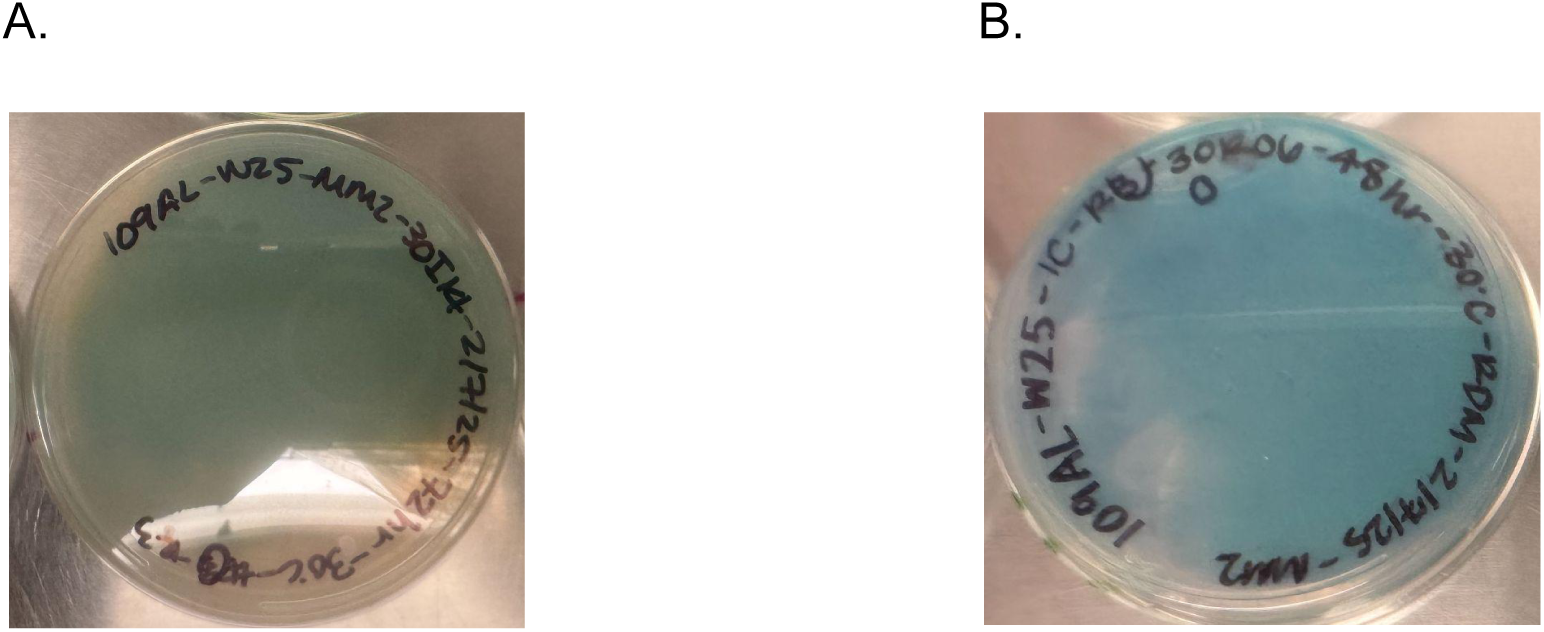
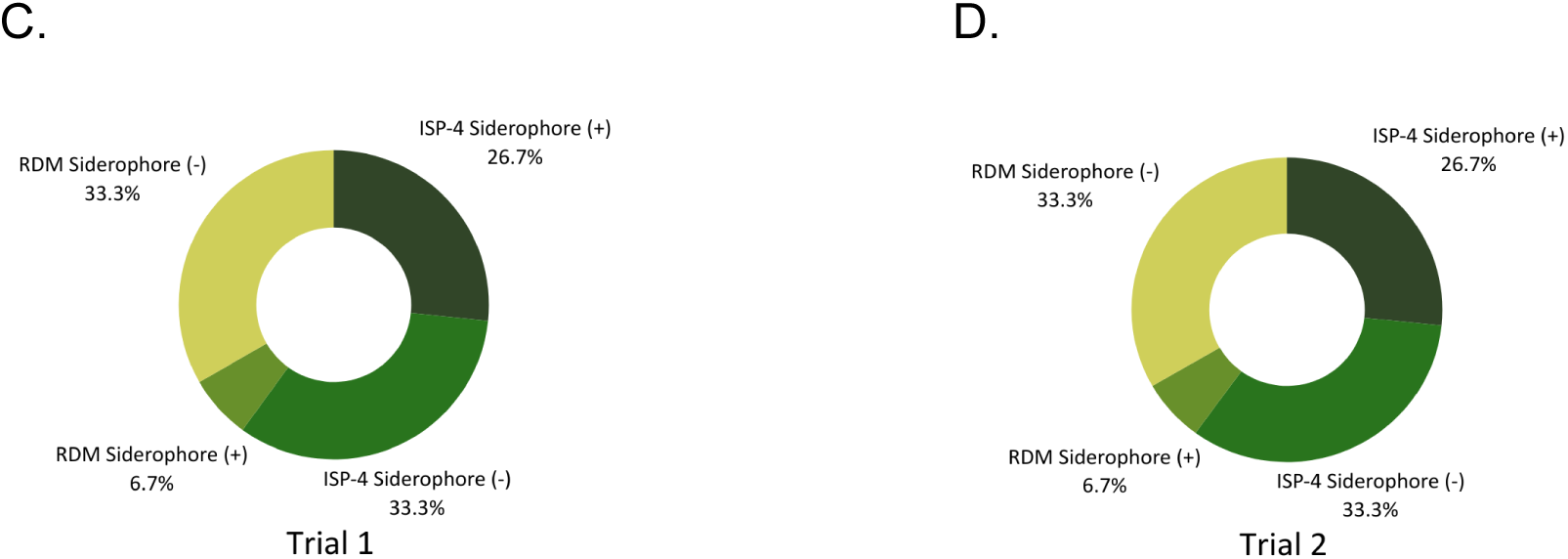
Siderophore Production of Isolates. **A)** A CAS overlay was poured over an ISP4 plate for isolate W25109CMM230I14. Color change from blue to blue and yellow was observed which demonstrated siderophore production. The negative control used for this experiment was *E. coli.* **B)** A CAS overlay was poured over an RDM plate for isolate W25109CMM230R06. There was no color change observed which demonstrated no siderophore production. **C)** After performing the siderophore production assay on all isolates, 33.4% of all isolates (grown on ISP4 or RDM) tested positive for siderophore production while 66.6% tested negative for Trial 1. **D)** After performing the siderophore production assay on all isolates, 33.4% of all isolates (grown on ISP4 or RDM) tested positive for siderophore production while 66.6% tested negative for Trial 2.

### Bacterial Effects on Carbohydrate Polymer Degradation

Carbohydrate polymer degradation assay assessed the metabolic capability of the microbes to break down complex carbohydrates such as, cellulose and xylan, into biofuels. A decrease in enzyme activity is seen after one, three, and eighteen days of a wildfire (Barreiro & DíazRaviña, 2021). However, the decomposed organisms created labile carbon available to the organisms. This is supported by **Figure 5A** and **5B**, showing the regions of hydrolysis by the pure isolates within the two trials. RDM, specifically isolate R20, showed the most distinct hydrolysis zone, as seen in **Figure 5C** and **5D**. These results are vital in understanding how droughts impact microbial activity in their ability to produce biofuels for the environment.

**Figure 3.**
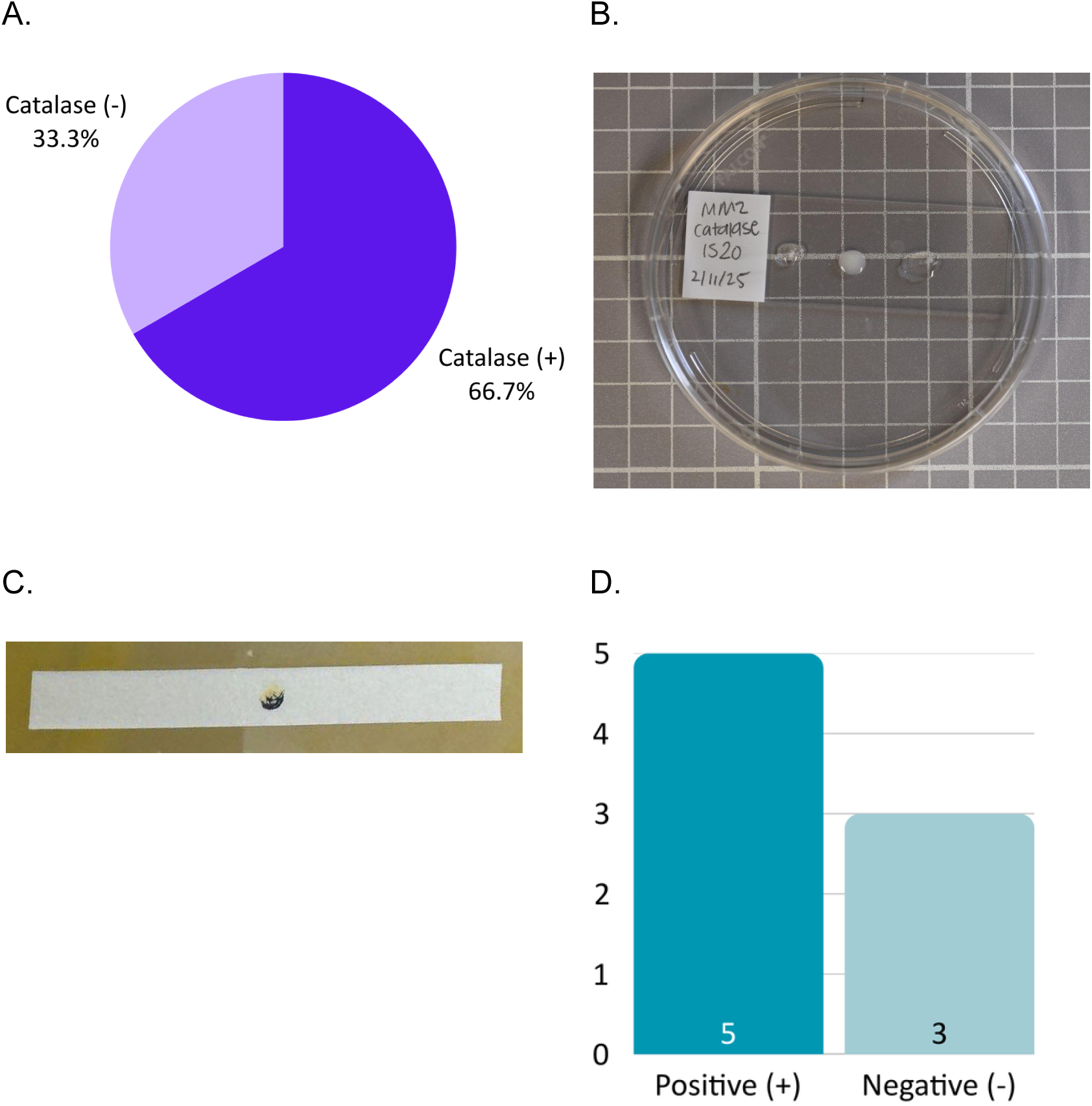
Catalase & Oxidase Production of Isolates. **A)** Of the six gram (+) isolates, 33.3% (2) did not produce catalase and 66.7% (4) produced catalase. **B)** A catalase test was performed on isolate W25109CMM230I20 in between catalase tests performed on negative (*Enterococcus*) and positive (*E. coli*) controls. Bubbles were produced upon the introduction of hydrogen peroxide, indicating the isolate produced catalase. The left side is the isolate, the middle is the positive control, and the right side is the negative control **C)** After performing the oxidase test on isolate W25109CMM230I02, the oxidase test paper showed a black or dark purple color change, indicating a positive result. d) After performing the oxidase test on all qualified isolates, 5 isolates tested positive for oxidase production and 3 isolates qtested negative.

**Figure 4.**
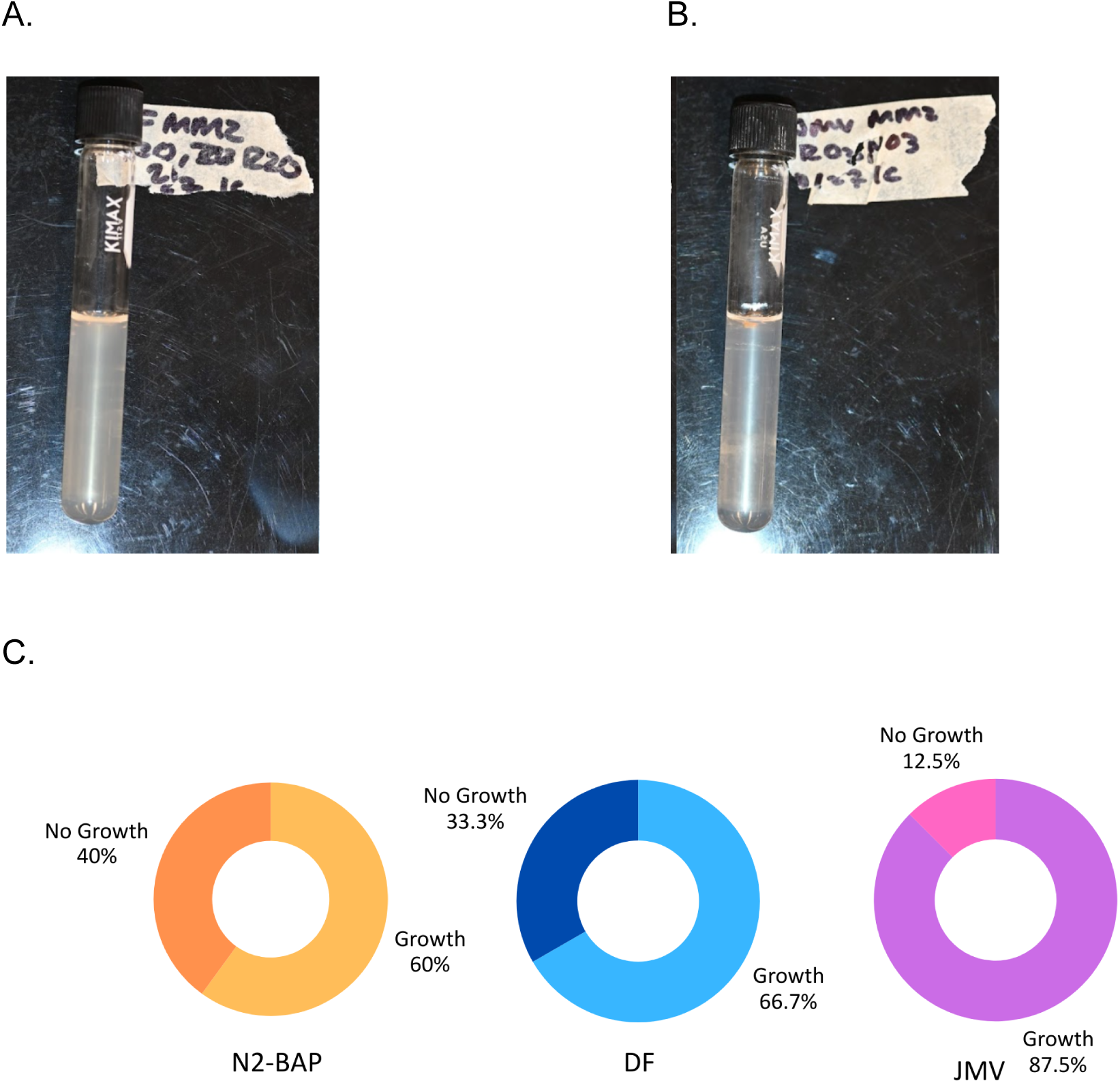
Nitrogen Fixation Results. **A)** Isolate W25109CMM230R20 was incubated on three types of media––N2-BAP, DF, and JMV. After incubation, it was found that W25109CMM230R20 only grew on N2-BAP and JMV. **B)** After incubation on the three types of media, it was found that isolate W25109CMM230R03 grew on all three types of media. **C)** After analysis of all isolates, it was determined that 60% of all purified isolates grew on N2-BAP, 66.7% grew on DF, and 87.5% grew on JMV.

**Figure 5.**
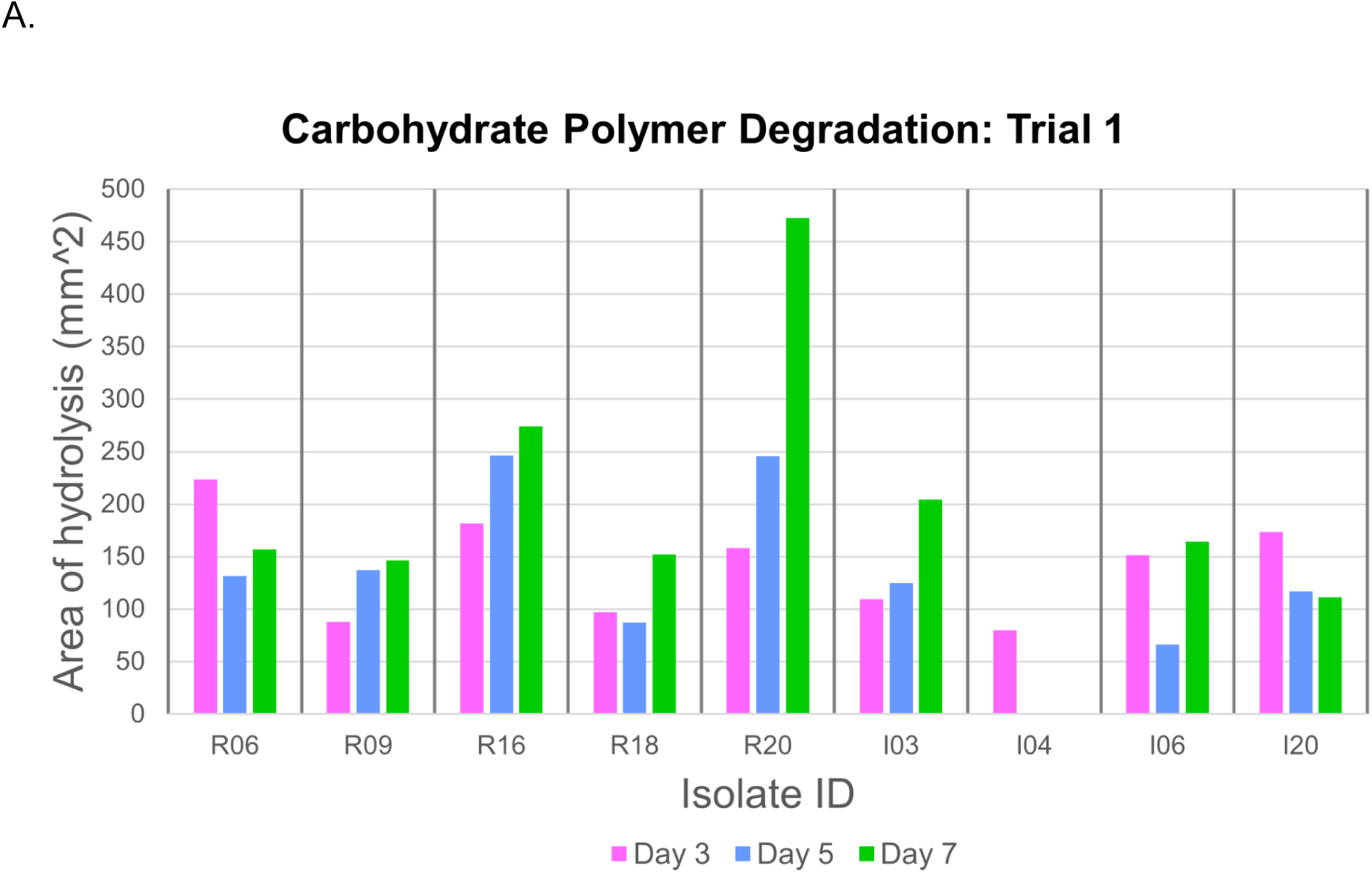
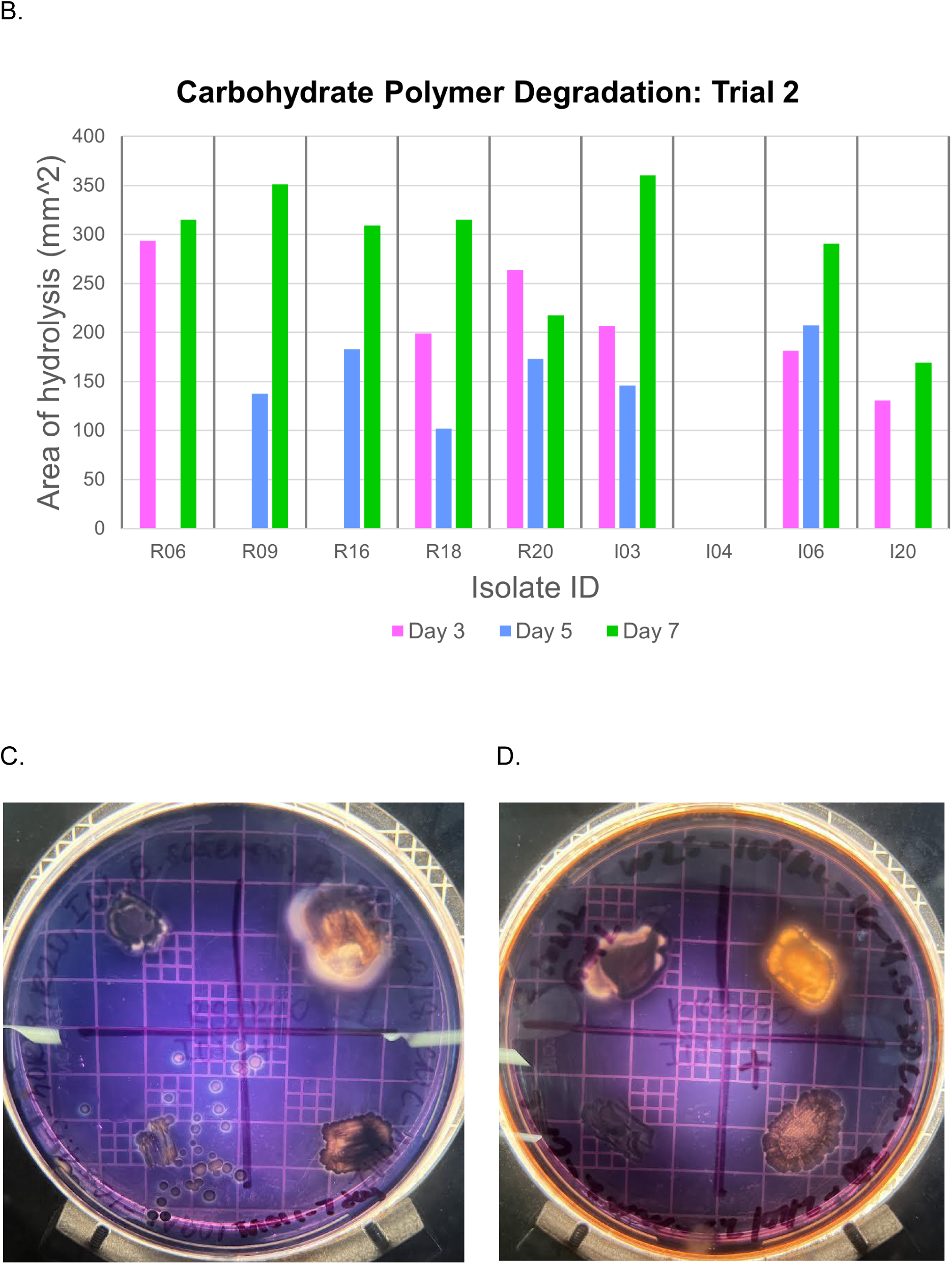
Carbohydrate Polymer Degradation Results. **A)** A graphical representation of Trial 1 of the carbohydrate polymer degradation assay. Zones of clearing were produced by the enzymes of each isolate after they were incubated for 3, 5, and 7 days on TY plates which contained diverse carbon sources. There was an increase in degradation with longer incubation periods which was depicted best by isolates cultured on RDM plates. **B)** A graphical representation of Trial 2 of carbohydrate polymer degradation assay. Zones of clearing were made by isolates capable of hydrolyzing the carbon rich plates. By day 7, the majority of the isolates with a positive result of this functional assay, showed an increase in the area degraded. **C)** Isolate W25109CMM230R20 showed a distinct zone of clearing formed by the bacteria’s enzymatic activity. After 7 days, it was able to hydrolyze the carbon sources on the TY plate, represented by the halo, which indicated the production of biofuels. **D)** R20 was the most clear result from all the isolates, including those tested from ISP4. It resulted in a glowing background where the bacteria was able to colonize and promote biofuel production.

### Identification of Nitrogen-Fixing Bacteria

Nitrogen fixation assesses the ability of the isolates to fix atmospheric nitrogen. Due to drought there was a decrease of mineralization in nitrogen cycles (Bogati & Walczak 2022). Of the 15 isolates plated on N2-BAP 60% or 9 (**Figure 4C**) of the isolates grew on the media. The 9 isolates were further analyzed on DF and JMV media which indicated that the bacteria were nitrogen fixators. In **Figure 4C**, 66.75% grown on the N2-BAP plates, cultivated on the DF selective media, confirming nitrogen fixation. JMV media, a nitrogen-free media, was used to see if the isolates potentially belong to the nitrogen-fixing *Burkholderia* family. **Figure 4C** shows 87.5% of the bacteria, grown on the N2-BAP plates, successfully developed on the JMV media. These significant outcomes indicated the percentage of nitrogen-fixing bacteria in the soil and their relation to the *Burkholderia* family.

## Biochemical Findings

### Determination of Catalase Activity

A catalase test was done on Gram(+) bacteria to check for the presence of catalase enzymes closely associated with aerobic respiration. Due to lower soil moisture there should be a decrease in bacterial enzymes (Bogati & Walczak 2022). As seen in **Figure 3B**, the left specimen depicts the isolate undergoing the chemical change, producing gas bubbles. In **Figure 3A**, 5 of the 6 isolates produced bubbles indicating the presence of catalase. These results are important as they may indicate that drought stressed soil bacteria may have an increase in aerobic activity.

### Effects of Oxidation

Oxidase tests were done on Gram (+) bacteria to check for the presence of cytochrome C, an electron transfer component utilized in aerobic respiration in the electron transfer chain for energy production. Due to lower soil moisture, aerobic respiration was reduced, decreasing cytochrome C activity (Schimel 2018). **Figure 3C** shows a positive reaction in which the oxidase paper appeared black/ dark purple; negative results show no color change. Of the 8 tested isolates, 5 are positive for oxidation, as shown in **Figure 3D**. These results indicate that there may be an increase of activity for cytochrome C.

## Metagenomic analysis

### Visual Representation of Bacterial Abundance

In dry soil, there should be an increase in drought-tolerant microbes. The relative abundance seems to shift in low moisture levels which includes a decrease in Gram-negative phyla Proteobacteria, Verrucomicrobia, and Bacteroidetes and increase in Gram-positive phyla Firmicutes and Actinobacteria (Naylor 2018). The Winter 2025 cohort displayed a large abundance of the most favored classes, *Actinobacteria*, *Alphaproteobacteria*, and *Bacilli* in **Figure 6A**, refuting the expected results since there is an increase in abundance of *Proteobacteria*. As seen in **Figure 6B**, the chart shows strong similarities between the two sites which demonstrates the relative abundance of the bacterial classes. Although the sites vary in specific quantity of each class by almost double, the proportions are relatively similar throughout the classes. **Figure 6C** displays the comparison between native and invasive plants of the Winter 25’ cohort at the class level. The most abundant classes include *Actinobacteria*, *Alphaproteobacteria* and *Bacilli* which are relatively the same in ratios. The percentage of class abundance becomes distinguished when comparing Betaproteobacteria between native and invasive plants. Overall, there are more similarities than differences between the groups and this could be due to environmental conditions such as the drought season and wildfires, allowing the most resilient bacteria to thrive in harsh conditions.

**Figure 6.**
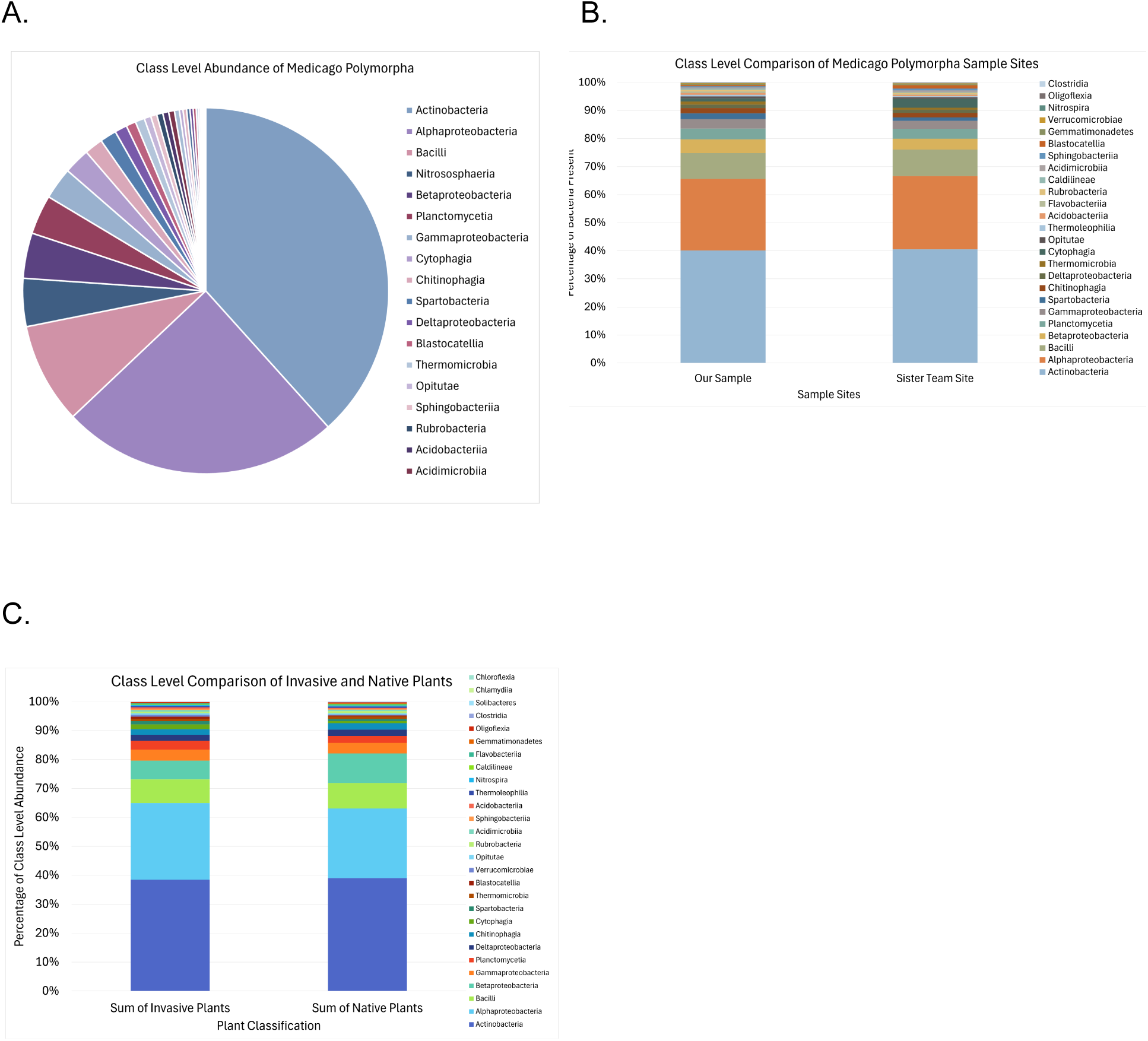
Class Level Analysis Using ASV Tables and Pivot Tables. **A)** Actinobacteria, Alphaproteobacteria, Bacilli are displayed as the dominant classes within the entire soil collection of *Medicago polymorpha* in the pie chart. **B)** The class comparison between the two soil samples includes ~40% belonging to Actinobacteria and ~20% being Alphaproteobacteria. Our sample site contained 1621 Actinobacteria and 1030 Alphaproteobacteria while the sister site contained 3314 and 2136, respectively. The quantity of bacteria present was a 1:2 ratio among the sample sites. **C)** The native and invasive classes were compared to see the relative abundance among Sage Hill. Actinobacteria, Alphaproteobacteria and Bacilli are the most common among the region. However, the invasive plants have a higher count by at least 1000 among the three most prevalent classes.

### Statistical Analysis of Metagenomic Profiles (STAMP)

The use of STAMP will help analyze the statistically significant variations and give a broader understanding of our data along with past and present cohort(s) data. These variations between taxonomic and functional profiles of Medicago would assess similarities and differences to native and invasive plants by comparing cohorts and by specific bacterial taxa. STAMP provides a graphical interface that permits exploration of statistical results and generation of publication-quality plots (Parks, 2014). **Figure 7A** provides a PCA plot on how Medicago is more of a native than invasive due to its similar bacterial species in native plants, indicating stress due to the drought. **Figure 7B** PCA plot provides bacterial classes from 2024 and 2025 cohorts, indicating Medicago had similar bacteria present in the wetter seasons. **Figure 7C**, PCA plot shows different water content as 2024 had an atmospheric river vs dryness conditions in 2025. In **Figure 7D**, the bar chart indicates how *Actinobacteria* is present more in extreme dryer and fire conditions than in wetter seasons. As for **Figure 7E**, an extended error bar plot has *Actinobacteria* and two classes of *Proteobacteria* shared between 2024 cohort and Medicago. Lastly, **Figure 7F** is a comparison of Medicago with the 2024 cohort to assess similarities present in both cohorts and showing *Actinobacteria* having the highest presence due to the drought conditions. These figures indicate that due to the drought-related conditions and wildfires a decrease in microbial activity Proteobacteria and Actinobacteria will be present, indicating that the rhizosphere of Medicago is affected by drought-related conditions.

**Figure 7.**
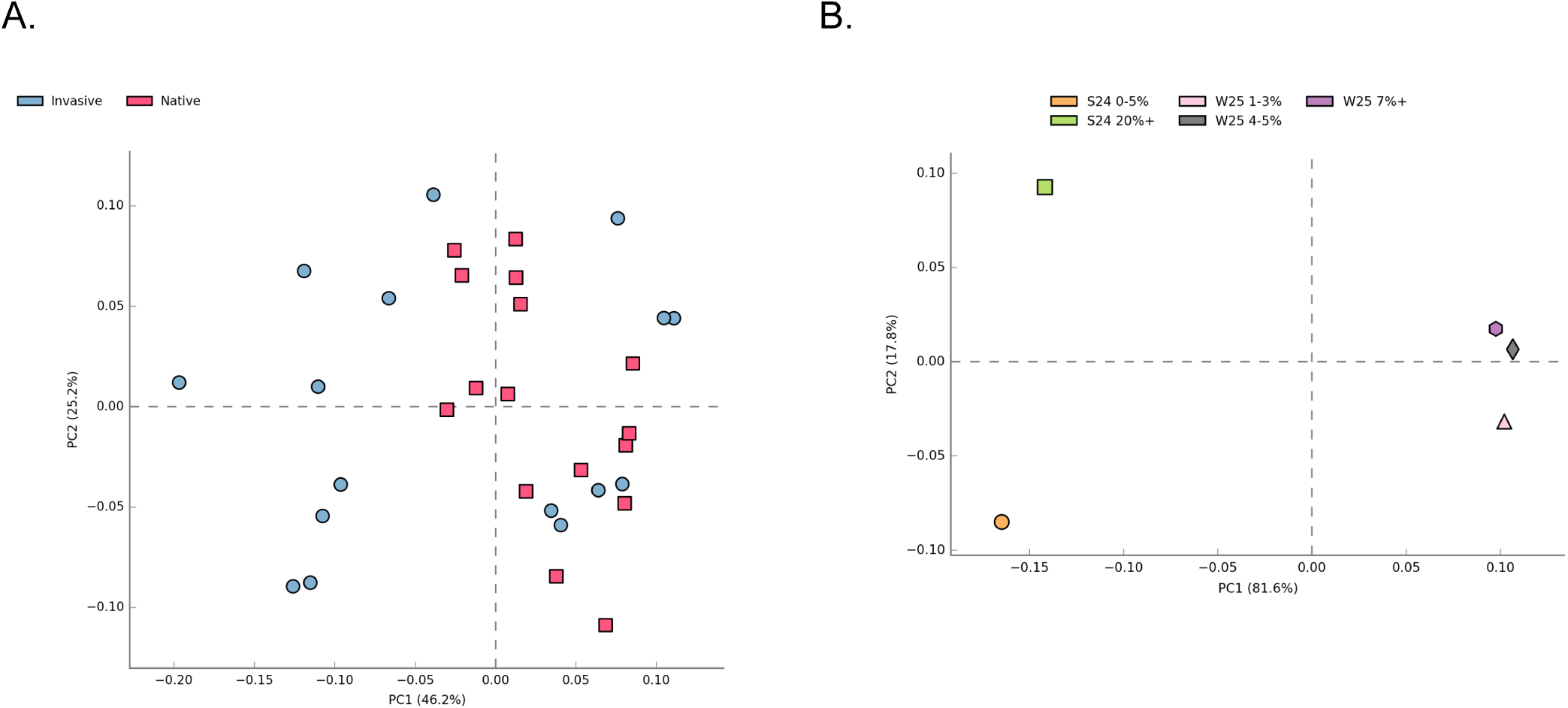
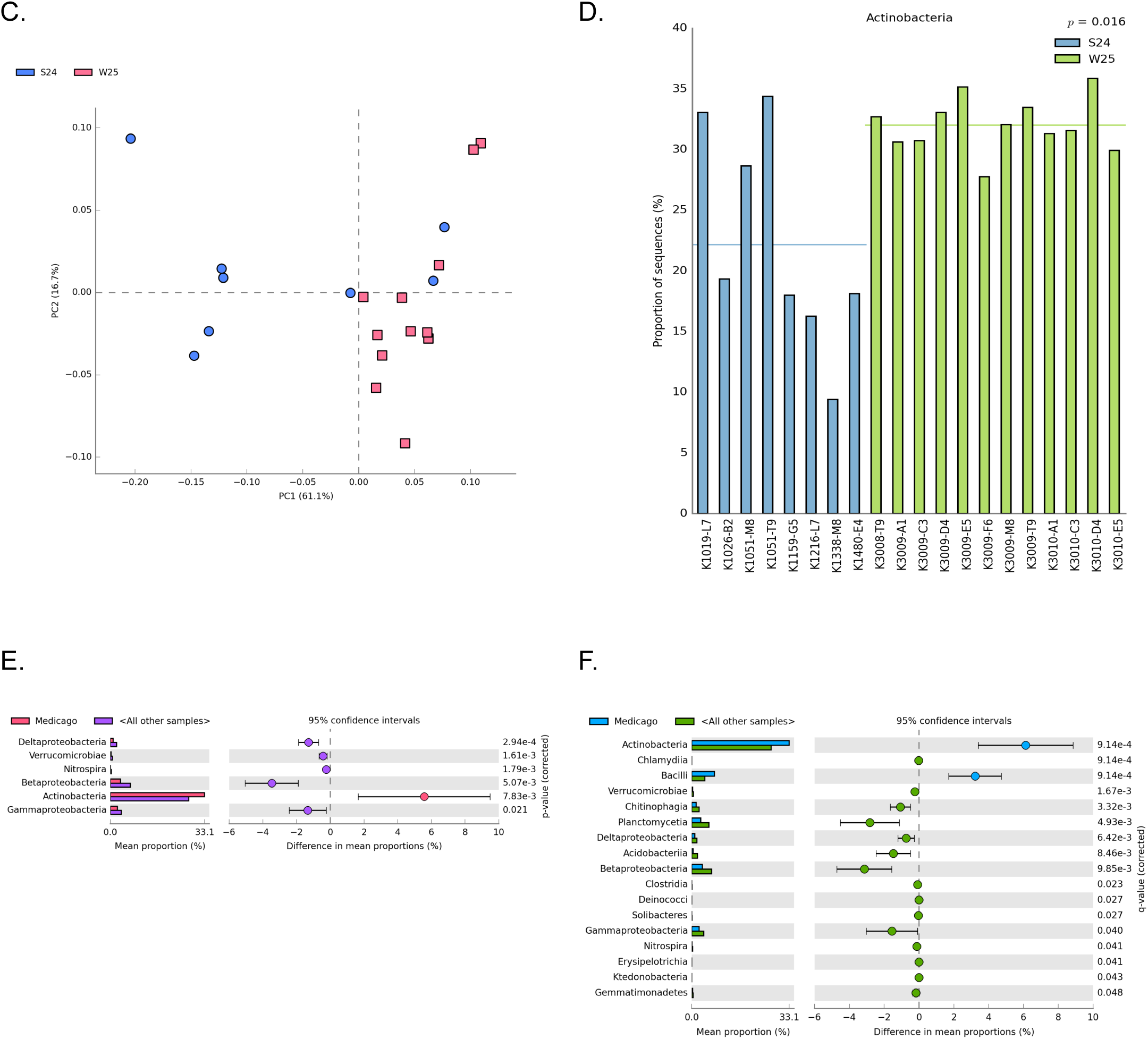
Comparing cohorts and bacterial lineages to determine statistical significance. **A)** The PCA plot represents the class level variation among soil site samples collected from Spring 2024, Winter 2024, Winter 2025 cohorts. This displays *Medicago* being closely related to native plants, based on the bacteria present. **B)** The PCA plot demonstrates the close relationship of bacterial classes from Spring 2024 and Winter 2025 where *Medicago* contains microbes closely related to the wet season. **C)** The soil moisture variations showed the difference in water content across the 2 cohorts, even though they contain similar microbial compositions. **D)** Spring 2024, the season consisting of the atmospheric river, showed lower levels of *Actinobacteria*. The Winter 2025 cohort had a statistically significant increase during the drought and wildfires, displaying their resilience to harsh climate conditions. **E)** The extended error bar plot expressed *Actinobacteria* and two classes of *Proteobacteria* being the shared prevalent classes amongst *Medicago* and Spring 2024 cohort. **F)** *Medicago* of Winter 2025 was compared to Winter 2024 as a standardized measurement to assess the relative similarities across the two cohorts, depicting *Actinobacteria* as the most similar class.

## Discussion

The primary goal of this study was to assess how drought impacts microbial composition and function in the rhizosphere of *Medicago*, an invasive legume species not previously examined in this context. We hypothesized that drought would suppress microbial activity and select for drought-tolerant taxa like *Actinobacteriota* and *Acidobacteria*. Overall, our results partially supported this hypothesis. Cultivation-dependent assays revealed a decrease in production of energetically expensive secondary metabolites (only 33.3% of culturable isolates are siderophore producers), but nitrogen fixation and carbohydrate degradation (both 60%) remained prevalent. This could suggest functional resilience within the rhizosphere under low soil moisture (4.4%). Cultivation-independent analysis revealed relatively increased abundance of *Actinobacteria* and *Firmicutes* per expectations, but unexpectedly high levels of *Alphaproteobacteria* partly refuted our expectations shaped by existing literature.

Unexpectedly, *Proteobacteria* remained abundant which contradicts published literature indicating Gram-negative taxa usually decline under drought conditions (Naylor & Coleman-Derr, 2018). This may be due to the atmospheric river event in the previous cohort, as rewetting pulses are known to favor fast-growing taxa (Schimel, 2018). *Acidobacteria* were also less abundant than expected, possibly due to their oligotrophic nature that was disrupted by low carbon availability during drought (Bogati & Walczak, 2022). The relative prevalence (60%) of nitrogen-fixing *Burkholderia* was also surprising. The presence of this could suggest that free-living diazotrophs may be favored in nutrient-depleted soils where nitrogen mineralization is reduced.

Functionally, the isolates that were capable of polymer degradation, especially RDM20, also challenged the initial idea that drought suppresses enzyme activity, instead suggesting that drought-tolerant taxa retain or upregulate pathways for accessing complex carbon sources (Barreiro & Díaz-Raviña, 2021). Regarding siderophore production, a proposed explanation would be that resource limitations as a result of drought constrains energetically costly activities (which siderophore production would be a part of) (Breitkreuz *et al*., 2021). These findings not only align with established patterns of increased *Actinobacteria* and *Firmicutes* under drought (Breitkreuz *et al*., 2021) but also suggest that plant-associated communities can maintain greater stability when it comes to environmental perturbations.

Despite differences in site moisture and plant identity, metagenomic analysis revealed that microbial class-level profiles remained similar across sites, possibly indicating that drought selects for rather than eliminates rhizosphere communities and that plants may buffer microbial communities against extreme stress or vice versa through plant-microbe relationships. Novel insights made during this study include the discovery of multiple free-living nitrogen fixers, suggesting that drought may promote mutualistic adaptations that are beyond the usual root nodule partnerships in plants like *Medicago*.

However, like most microbiology-based studies, our conclusions are constrained by cultivation bias and sequencing limitations. Only culturable microbes were tested for function which likely heavily underestimated community diversity. 16S data lacked sufficient resolution for genus and species-level dynamics and did not account for functional genes, again, limiting the extent of our conclusions. In addition, potential misidentification of isolates due to limited databases and visual confusion from pigmentation in the siderophore assays may also skew interpretation.

Despite these limitations, the study offers practical relevance. The persistence of nitrogen-fixing and degradative traits implies that drought-adapted microbes could support sustainable agriculture. Isolates with these traits might be developed into microbial soil additives to enhance crop resilience without reliance on fertilizers. Identifying microbial traits like catalase, oxidase, and polymer degradation in drought-tolerant microbes could provide targets for future microbial screening aimed at stress resilience, especially in communities commonly impacted by drought.

A short-term experiment would be to grow *Medicago* and a native comparison plant) in soil under controlled watering vs. drought conditions. By sampling the rhizosphere soil at multiple time points (e.g. before drought, during drought, and after rewetting), one could track microbial community dynamics through the onset and relief of drought, tightly controlling these environmental perturbations. Successful high-throughput 16S rRNA gene sequencing would be used to provide statistically sound data on how each major taxon changes over time. Coupling this with measurements of soil moisture would more robustly link microbial changes to plant condition.

Our existing culture collection could be expanded and leveraged in future studies by attempting to isolate a broader diversity of microbes from drought-impacted soils, including those that might have been missed by using longer incubation times, different media, etc.. Once isolated, strains of interest (specifically those that are dominant in sequencing data or with promising traits, like strongest nitrogen fixers or polymer degraders) would be characterized in detail. This involves identifying them via full-length 16S or whole-genome sequencing and testing their physiological limits (such as tolerance to desiccation and/or high osmotic pressure).

On a broader scale, a beneficial future direction could be to establish a long-term field study at Sage Hill to monitor microbial community resilience over multiple seasons and years. This could involve periodic soil sampling and analyzing not just DNA but also soil enzyme activities and nutrient level changes. In addition, including fungal community analysis would fill a gap in our current study, as fungi play major roles in drought, often increasing in relative importance as bacteria decline (de Vries & Shade, 2013).

To further overcome cultivation biases and study interactions, techniques like iChip could help grow previously unculturable drought survivors by simulating their natural environment. Constructing synthetic communities in the lab, consisting of a defined mix of isolates from our collection (potentially a combination of *Actinobacteria*, *Firmicutes*, and *Proteobacteria* representative of the drought community of *Medicago*), could also allow for controlled experiments on community function. We could vary water availability in vitro and measure outputs like carbon concentration or plant seedling growth promotion to directly assess how interactions among these microbes respond to drought which would deepen our understanding of microbial community composition changes under stress.

## Acknowledgements

We would like to sincerely thank Shaun Liu, Tara Mata, and Aaron Holaway for their guidance and expertise with our bioinformatical analyses and wet-lab procedures. CALeDNA assisted in DNA analyses for our cultivation independent methods. Thank you to the Dean of the Life Sciences Division and the Teaching and Learning Center Seed Grant for the funding of our project.

## Author Contributions

Soil enrichment and cultivation by serial dilutions were completed by N.O., B.P., S.R., and R.M. N.O., B.P., and S.R. performed the purification rounds. S.R. maintained streak plating with R.M. contributing to one week of maintenance. Gram staining was performed by all members. Manual checking for purity was performed by S.R., N.O., and R.M.; photos taken by R.M. Soil pH test, determination of moisture content, and determination of nitrogen content were performed by R.M. Determination of active carbon, potassium, and phosphorus content were performed by N.O. and R.M. Testing of nitrogen fixation capabilities was performed by S.R and B.P. Carbohydrate polymer degradation was performed by S.R. with first trial inoculation performed by N.O. CAS overlays were performed by N.O. and B.P. Catalase and oxidase test was performed by B.P. and N.O. Pivot tables generated by S.R. STAMP figures generated by S.R. with contributions from N.O and R.M. N.O., B.P., S.R., and R.M. drafted paper. S.R., N.O. and B.P. formatted and revised paper.T.B. supervised overall research.

